# Comparison and benchmark of long-read based structural variant detection strategies

**DOI:** 10.1101/2022.08.09.503274

**Authors:** Jiadong Lin, Peng Jia, Songbo Wang, Kai Ye

## Abstract

**Background:** Recent advances in long-read callers and assembly methods have greatly facilitated structural variants (SV) detection via read-based and assembly-based detection strategies. However, the lack of comparison studies, especially for SVs at complex genomic regions, complicates the selection of proper detection strategy for ever-increasing demand of SV analysis.

**Results:** In this study, we compared the two most widely-used strategies with six long-read datasets of HG002 genome and benchmarked them with well curated SVs at genomic regions of different complexity. First of all, our results suggest that SVs detected by assembly-based strategy are slightly affected by assemblers on HiFi datasets, especially for its breakpoint identity. Comparably, though read-based strategy is more versatile to different sequencing settings, aligners greatly affect SV breakpoints and type. Furthermore, our comparison reveals that 70% of the assembly-based calls are also detectable by read-based strategy and it even reaches 90% for SVs at high confident regions. While 60% of the assembly-based calls that are totally missed by read-based callers is largely due to the challenges of clustering ambiguous SV signature reads. Lastly, benchmarking with SVs at complex genomic regions, our results show that assembly-based approach outperforms read-based calling with at least 20X coverage, while read-based strategy could achieve 90% recall even with 5X coverage.

**Conclusions:** Taken together, with sufficient sequencing coverage, assembly-based strategy is able to detect SVs more consistently than read-based strategy under different settings. However, read-based strategy could detect SVs at complex regions with high sensitivity and specificity but low coverage, thereby suggesting its great potential in clinical application.

## Background

Structural variants (SVs) comprise different subclasses, such as deletions, insertions, etc, are playing important roles in both healthy and disease genomes. To date, researchers have made great progress in discovering and genotyping SVs in diverse populations with short-read data, but SVs at repetitive regions remain challenging due to limited read length [1]. Even in non-repetitive regions, SVs such as insertions are missed by approaches relying solely on short-reads [2]. Single-molecule sequencing (SMS) technologies, such as Pacific Bioscience (PacBio) and Oxford Nanopore Technology (ONT), have emerged as superior to short-read sequencing for SV detection and thus revealing a number of novel functional impact of SVs missed by short-read data [3, 4]. Long reads also improved SV detection in genetic diseases [5-7] and cancers [8-14] where SVs are usually undetectable or misinterpreted by short-read, such as the ONT data reveals 10,000bp Alzheimer’s disease associated *ABCA7* Variable Number Tandem Repeats (VNTR) expansion that are missed by short-read data [15]. The outstanding detection performance and the great demand of long-read based applications raises a problem of selecting proper strategy for SV detection. For example, the Chinese [16] and Icelander [17] cohort studies detect SVs directly from reads alignment. Another clinical study showed a likely pathogenic SV can be identified from reads eight hours after enrollment, while similar results were received two weeks using traditional diagnose approaches [18]. Instead of detecting directly from reads, the advances in assembly methods promote SV detection from haplotype-aware assemblies, such as the study conducted by Human Genome Structural Variation (HGSV) consortia, revealed 107,590 SVs with HiFi assemblies, of which 68% are not discovered by short-read sequencing [3, 19].

Currently, almost all long-read based studies use either read-based strategy (i.e., detecting directly from read alignment) or assembly-based strategy (i.e., detecting from alignments of de novo assemblies) for SV detection. The assembly-based strategy requires an extra step for haplotype-aware assembly, but the following steps of the two strategies are similar and usually contains two parts. Firstly, the variant signatures are identified and gathered from two types of aberrant alignments: intra-read and inter-read. Intra-read alignments are derived from reads spanning the entire SV locus, resulting deletion and insertion signatures. Inter-read alignments are usually obtained from the supplementary alignments and SV signatures could be identified from inconsistencies in orientation, location and size during mapping, from which translocation as well as large deletion, duplication and inversion signatures are identified. Secondly, callers typically cluster and merge similar signatures from multiple aberrant alignments, delineating proximal signatures that support putative SV. Nearly all read-based callers developed in the past five years, such as Sniffles [20], pbsv, cuteSV [21], SVIM [22], NanoVar [23], NanoSV [24], and Picky [9], detect SVs through combinations of signatures obtained from inter-read and intra-read alignments but differ in their signature clustering heuristics. While different from the above methods, SVision applied a deep-learning approach to directly recognize different SV types from the variant signature sequences. As for assembly-based callers, such as Phased Assembly Variant (PAV) and SVIM-ASM [22] use the alignment of whole genome assembly as input, from which aberrant inter-contig and intra-contig alignments are collected and used for SV detection. Most importantly, accumulating studies have claimed that the assembly-based detection strategy is able to comprehensive detect SVs and characterize non-templated insertions [1, 3, 19]. Though a number of studies have demonstrated the advances of using long-read toward short-read data, it lacks systematic comparison of read-based and assembly-based strategy. Therefore, to help users, it is important to quantitatively assess and compare the stability and usability of the two strategies, especially for SVs at complex genomic regions. Moreover, the potential weakness of different strategies needs to be investigated, so that new developments in the field could focus on improving current methods.

In this study, a widely-used benchmark material, HG002 genome, was selected to compare and benchmark the two strategies. Moreover, according to methods reviewed by a recent study [25], we selected four read aligners, two assemblers for HiFi datasets, two assemblers for ONT datasets, one contig aligner, one phasing algorithm, five read-based callers and two assembly-based callers (**Methods**). We then evaluated the impact of detection settings (i.e., aligners and assemblers) and sequencing settings (i.e., read length, sequencer and coverage) on both strategies. Briefly, the impact of sequencing settings was first assessed for each strategy across all datasets based on datasets concordant and unique SVs, and the detection and sequencing settings affected strategy concordant SVs were further assessed (**Fig. 1a**). Additionally, the impact of detection settings on each strategy were examined on each dataset based on aligner concordant and assembler concordant SVs (**Fig. 1b**). For concordant SVs, we also assessed their breakpoint difference, where the breakpoint standard deviation (BSD) smaller than 10bp were classified as breakpoint accurately reproduced concordant SVs (**Fig. 1c**). Furthermore, for both strategies, their recall and precision of detecting well curated SVs, especially those at challenging medically relevant autosomal genes (CMRG), were assessed and cross-compared under different sequencing settings.

**Fig. 1.**
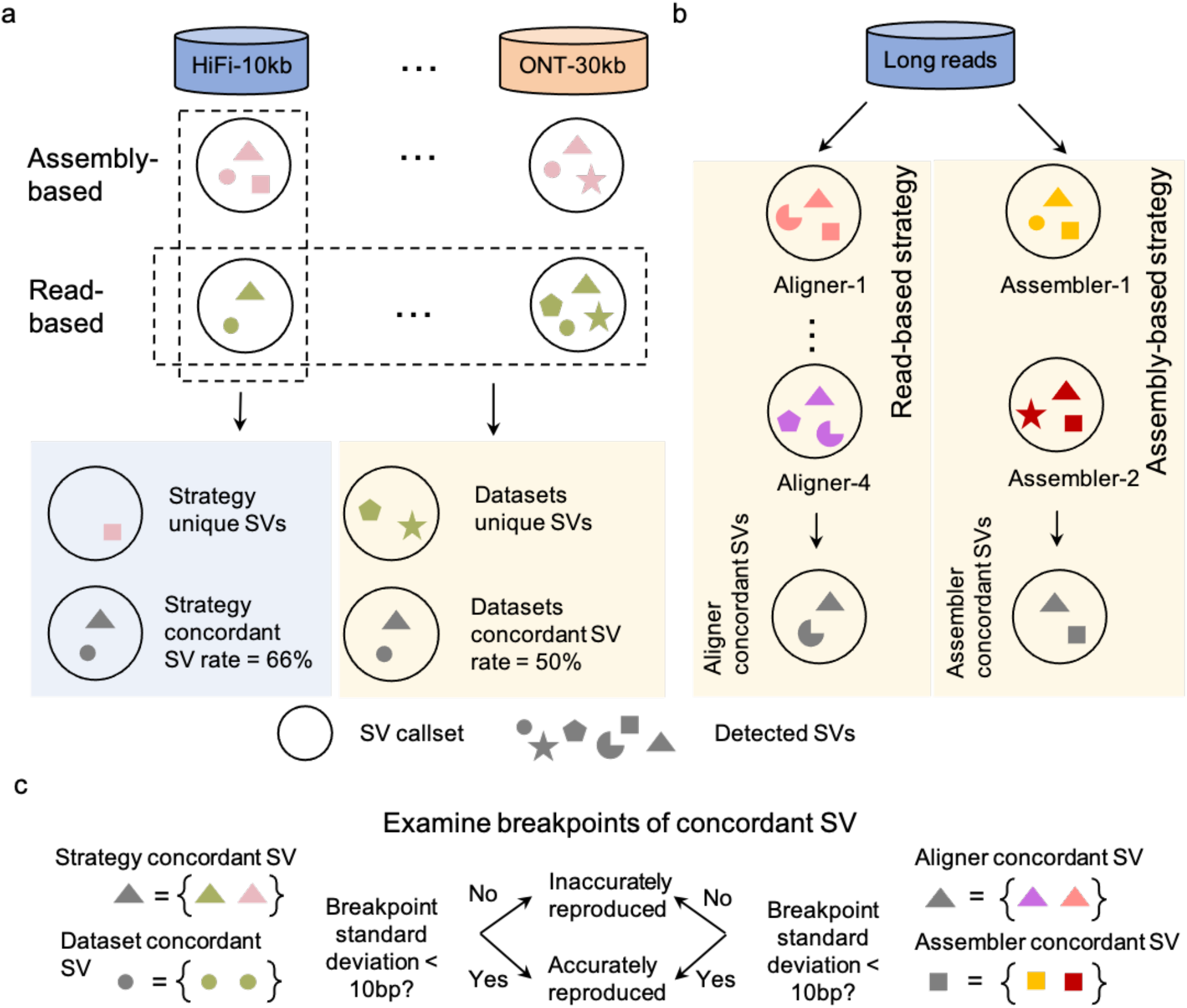
Schematic summaries of assessing the impact of different settings on each strategy and between strategies. **a**. Examining the impact of sequencing settings on each strategy based on datasets unique and concordant structural variants (SVs). Moreover, the impact of detection settings on strategy concordant SVs was assessed on each dataset. **b**. For each strategy, the impact of detection settings, i.e., aligners and assemblers, was assessed on each dataset based on aligner concordant SVs and assembler concordant SVs. **c**. Examining the breakpoint difference of concordant SVs, where the breakpoint standard deviation of concordant SVs smaller than 10bp was classified as breakpoint accurately reproduced SVs, otherwise, it was termed as breakpoint inaccurately reproduced SVs.

## Results

### Impact of sequencing settings on each strategy

We totally generated 120 read-based callsets and 24 assembly-based callsets, while SVs at centromere and low mapping quality regions were excluded in the analysis (**Method**). Overall, assembly-based and read-based strategy detected a median of 20,827 and 23,611 SVs from HiFi datasets, respectively, while more SVs were detected from ONT datasets, i.e., a median of 22,009 for read-based and 29,162 for assembly-based (**Fig. 2a**). As expected, the SV size peaks for both strategies were observed at 300bp and 6,000bp, indicating SINE and LINE, respectively (**Supplementary Fig. 1a**). Moreover, the majority of the SVs (75%) were located at repetitive regions without sequencing platform bias, while two strategies differed at Simple Repeats regions consisted of either VNTR or short tandem repeats (STR) (**Fig. 2b**). As for SV types, assembly-based strategy detected more insertions than read-based callers due to longer sequence length (**Fig. 2c**). While read-based caller SVision detected comparable percentage of insertions as assembly-based strategy when detected from minimap2 or winnowmap aligned ONT reads (**Fig. 2c**). On the contrary, pbsv paired with ngmlr resulted in the fewest percentage of insertions among all six datasets (**Fig. 2c**). Additionally, different from assembly-based strategy, read-based callers also identified other SV types, such as duplication and even complex types (**Supplementary Fig. 1b**).

**Fig. 2.**
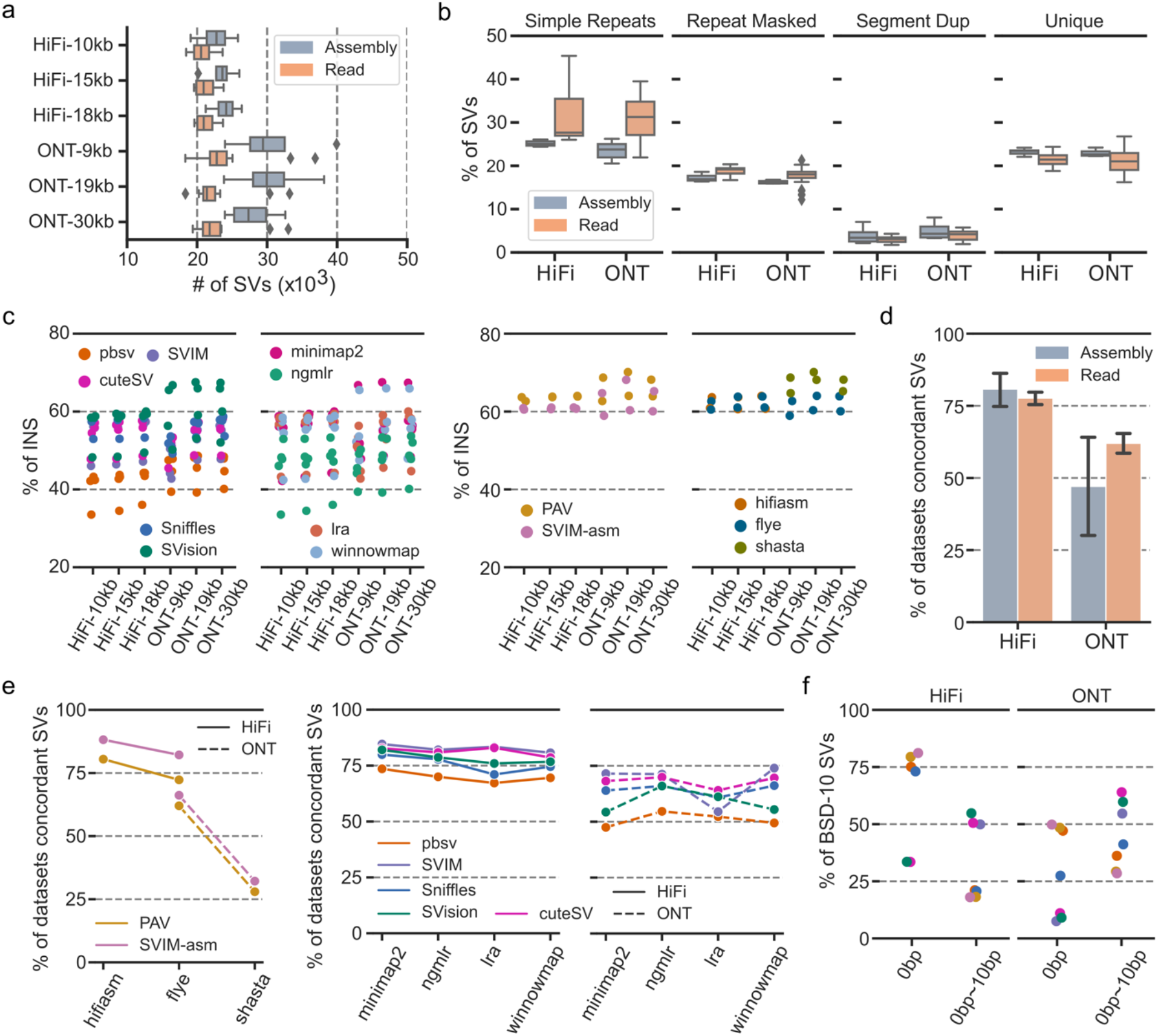
Summaries of the impact of sequencing settings on each strategy. **a**. The number of structural variants (SVs) detected by each strategy among datasets. **b**. The distributions of detected SVs among different genomic regions. **c**. The percentage of insertions affected by callers, aligners and assemblers. **d**. The percentage of dataset concordant SVs detected from HiFi and ONT datasets of each strategy. **e**. The percentage of dataset concordant SVs affected by callers, aligners and assemblers on HiFi and ONT datasets. **f**. The percentage of breakpoint accurately reproduced SVs (i.e., BSD-10 SVs) on HiFi and ONT datasets.

For each strategy, we further assessed the number and breakpoint of dataset concordant SVs. On average, detecting from HiFi reads, 75% and 80% of the dataset concordant SVs were identified for read-based and assembly-based strategy, respectively. However, the average dataset concordant SV rate of read-based strategy was higher than assembly-based strategy on ONT datasets, suggesting that read-based strategy was more versatile to different datasets (**Fig. 2d, Supplementary Fig. 1c**). Moreover, large variance of concordant SVs rate observed in ONT datasets suggested a great assembler bias, i.e., the average dataset concordant SV rate was 26% for shasta and it was 45% lower than detecting from assemblies created by flye (**Fig. 2e**). Comparably, as a critical setting for read-based strategy, the percentage of reproducible SVs detected from ONT reads was less affected by aligners when compared to assemblers did on assembly-based callers, i.e., the average dataset concordant SV rate for each aligner ranged from 50% to 75% (**Fig. 2e**). Furthermore, the average percentage of breakpoint accurately reproduced SV (i.e., BSD smaller than 10bp, BSD-10) on HiFi datasets was around 20% higher than that of ONT datasets (**Fig. 2f, Supplementary Fig. 1d**). For breakpoint inaccurately reproduced SVs, 65% (HiFi datasets) and 50% (ONT datasets) of them overlapped with simple repeat regions, while 5% of these SVs detected from ONT reads were found at segment duplication regions for both strategies (**Supplementary Fig. 1e**). We further investigated the impact of genomic regions on breakpoint accuracy and found that assembly-based strategy was able to detect more BSD-0 (i.e., BSD equals 0bp) SVs than read-based strategy, especially at simple repeat regions (**Supplementary Fig. 2>**). The above results showed that both strategies might overcall on ONT datasets and large variance of SVs at simple repeat regions was observed in read-based callsets. Though both strategies were able to detect SVs consistently from HiFi reads in terms of the concordant SV rate and their breakpoint consistency, the breakpoint of assembly-based calls were more accurate than read-based ones.

### Impact of aligners and assemblers on reproducible SVs for each strategy

Next, we examined the impact of detection settings (i.e., aligner for read-based and assembler for assembly-based) on each strategy (**Method**). For read-based strategy, around 50% of the SVs were detectable from all four aligners mapped reads, referring to as aligner concordant calls, while 30% of the SVs were only detected from one of the aligners and considered as aligner unique calls (**Fig. 3a, Supplementary Fig. 3-4**). The majority (80%) of the aligner concordant calls were found to be BSD-10 on both HiFi and ONT datasets (**Fig. 3b**). Notably, for pbsv, 75% of the aligner concordant calls’ breakpoints were BSD-0, which was 60% higher than other read-based callers, indicating that pbsv detected SV breakpoints were less affected by aligners than others, especially on HiFi datasets (**Fig. 3b**). As for assembly-based callers, 75% and 50% of the SVs were detectable from HiFi and ONT assemblies generated by two assemblers, respectively, and we termed these SVs as assembler concordant SVs (**Fig. 3c, Supplementary Fig. 5**). Remarkably, calling from HiFi reads, BSD-0 SVs took 98% of the assembler concordant SVs (**Fig. 3d**), which was 13% higher than pbsv and much higher than other read-base callers (**Fig. 3b**). Though the percentage of BSD-0 SVs detected from ONT assemblies was not comparable to HiFi assemblies, i.e., 60% for ONT and 98% for HiFi, assembly-based strategy was less affected by assemblers than that of aligners on read-based strategy. Moreover, we noticed that the percentage of BSD-0 aligner and assembler concordant SVs increased as the read length increasing (**Fig. 3b, Fig. 3d**). This might due to the Guppy version used for ONT base calling (**Supplementary Table S1**).

**Fig. 3.**
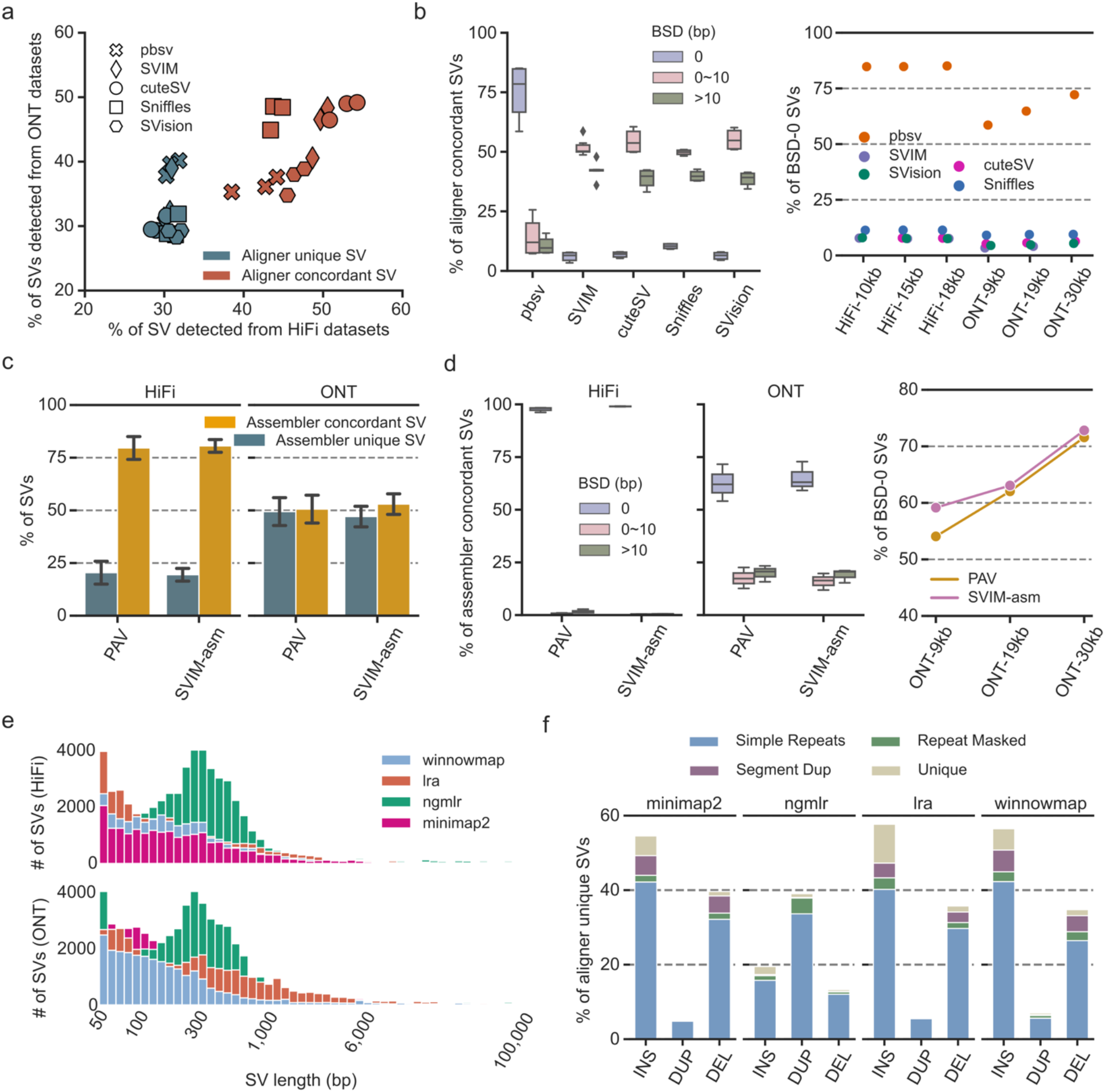
Summaries of the impact of detection settings on each strategy. **a**. The percentage of aligner unique and aligner concordant structural variants (SVs) detected from HiFi (*x*-axis) and ONT (*y*-axis) datasets. **b**. The percentage of breakpoint accurately reproduced SVs (i.e., BSD-10 SVs, right panel) and breakpoint identically reproduced SVs (i.e., BSD-0 SVs, left panel) identified from read-based callsets. **c**. The percentage of assembler unique and concordant SVs detected from HiFi and ONT datasets. **d**. The percentage of breakpoint accurately reproduced SVs (i.e., BSD-10 SVs, right panel) and breakpoint identically reproduced SVs (i.e., BSD-0 SVs, left panel) identified from assembly-based callsets. **e**. The size distribution of aligner unique SVs. **f**. The SV types among aligner unique SVs at different genomic regions.

**Fig. 4.**
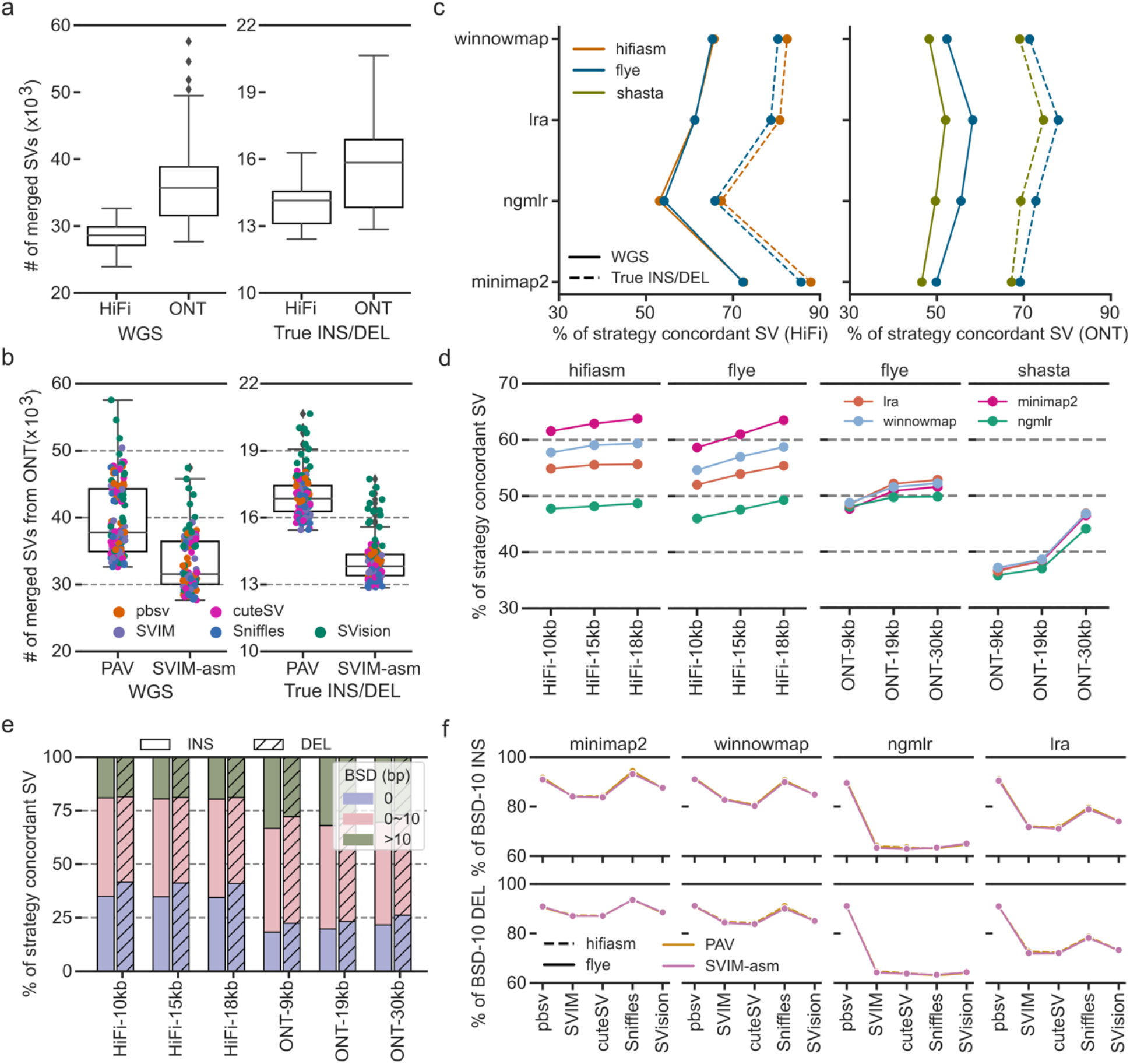
Summary of impact of detection and sequencing settings on the strategy concordant structural variants. **a**. The number of structural variants (SVs) in the nonredundant callset merged from read-based calls and assembly-based calls at whole genome scale (WGS) and true INS/DEL regions. **b**. The number of structural variants (SVs) in the nonredundant callset merged from read-based calls and assembly-based calls detected from ONT reads at WGS and true INS/DEL regions. **c**. The average percentage of strategy concordant SVs affected by assembler and aligner pairs at WGS and true INS/DEL regions. **d**. The average percentage of strategy concordant SVs on each dataset. **e**. The percentage of concordant SVs of different breakpoint standard deviation among datasets. ‘0’, breakpoint standard deviation equals 0bp. ‘0∼10’, breakpoint standard deviation large than 0bp but smaller or equal to 10bp. ‘>10’, breakpoint standard deviation large than 10bp. **f**. The percentage of breakpoint accurately reproduced SVs (i.e., BSD-10 SVs) affected by aligner, assembler and callers evaluated on HiFi-18kb dataset.

In addition, most of aligner or assembler unique SVs were located at Simple Repeat regions (**Supplementary Fig. 6a**). Using these uniquely detected SVs, we were able to investigate the impact of aligners and assemblers on the SV size and types. For aligner unique SVs, a median of 2,151 SVs and 2,677 SVs were found in HiFi and ONT datasets, respectively (**Supplementary Fig. 6b**). However, 2.5 times more SVs, ranging from 100bp to 1,000bp, were uniquely detected from ngmlr aligned reads without platform bias (**Fig. 3e**). Moreover, a significant peak at 300bp was only observed for SVs detected from ngmlr aligned reads (**Fig. 3e**). In terms of SV types, around 17%, 39%, 38% and 33% of the unique calls were deletions detected from ngmlr, minimap2, lra and winnowmap alignments, respectively (**Fig. 3f**). Besides the bias for deletions, 37% of the ngmlr unique calls were duplications and it was around 30% higher than the average of other aligners. Additionally, the percentage of ngmlr unique insertions was 23%, but the average percentage was 46% for other aligners, suggesting that ngmlr preferred to generate duplication like alignment signature for read-based callers (**Fig. 3f**). We reasoned that this aligner bias was largely due to the mapping strategy adopted by ngmlr, where it splits read into non-overlapping 256bp sub-reads and maps them independently of each other [20]. Thus, a size peak was observed close to 300bp and insertions could be aligned as duplications where two sub-reads overlapped on reference genome. For assembly-based strategy, a median of 2,482 SVs and 7,976 SVs were identified from HiFi and ONT assemblies, respectively (**Supplementary Fig. 6c**). The size of SVs detected from hifiasm assembled HiFi contigs was enriched at 300bp, and most of SVs detected from shasta created ONT assemblies ranged from 50bp to 300bp (**Supplementary Fig. 6d**). We only observed the insertion bias among the assembler unique SVs, where around 78% of shasta unique SVs were insertions and most of these insertions were smaller than 300bp (**Supplementary Fig. 6e**). Taken together, the above results suggested that read-based calls, including their breakpoints, types and sizes, were greatly affected by aligners, while up to 80% of the SVs, consisting of 98% BSD-0 SVs, were detectable from HiFi assemblies created by different assemblers.

**Fig. 5.**
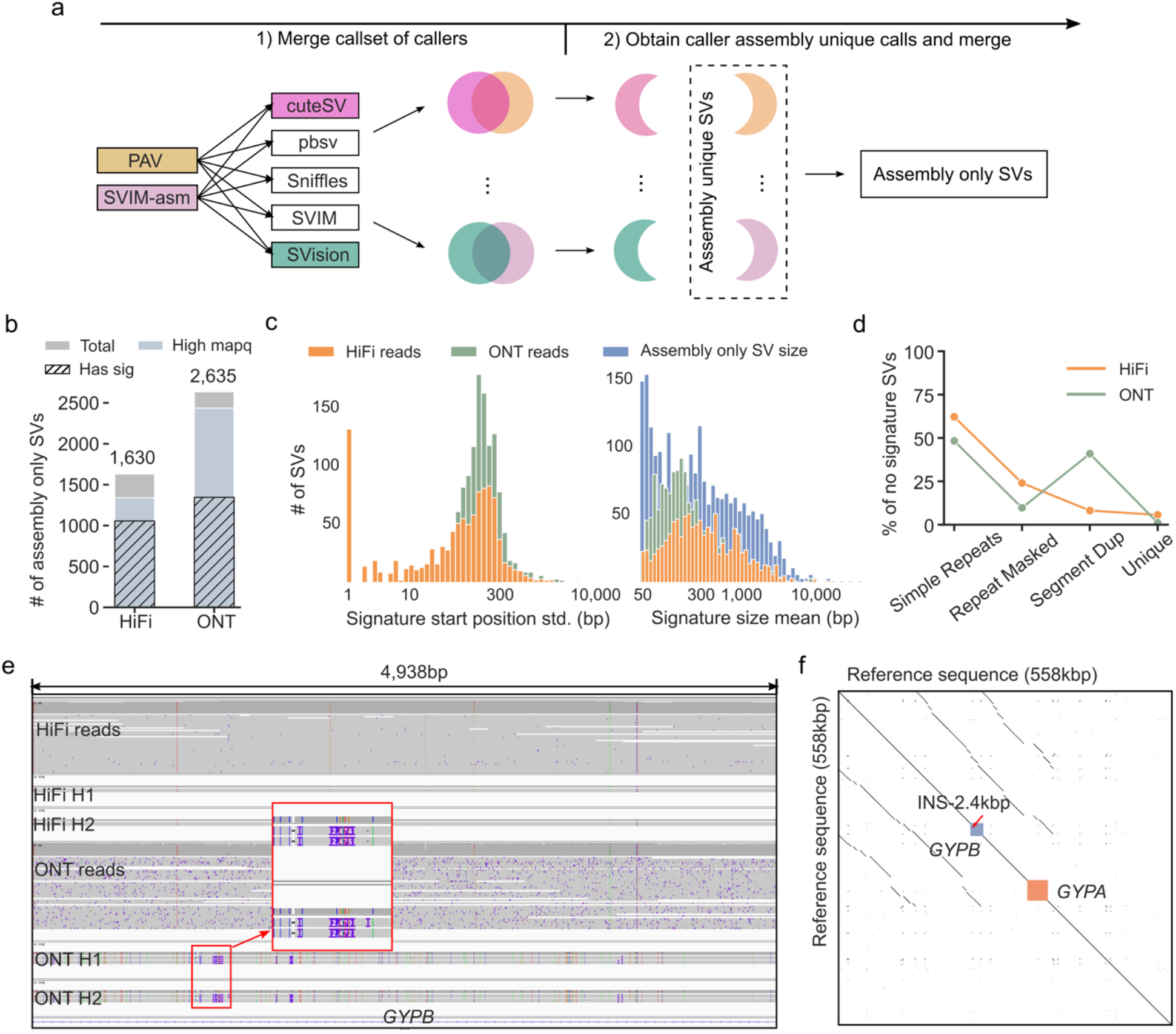
Examining assembly only structural variants. **a**. The schematic of obtaining assembly only structural variants (SVs) from assembly unique SVs. **b**. The number of all assembly only SVs, assembly only SVs at high mapping quality regions and assembly only SV loci containing at least five SV signature reads. **c**. The SV signature reads start position standard deviation (std) and the average length of identified signatures. **d**. The genomic region distribution of assembly only SVs without enough SV signature reads (smaller than five). **e**. The IGV alignment view of a 2.4kbp insertion incorrectly detected from ONT assemblies. **f**. The sequence Dotplot of local genome containing the insertional breakpoint shown in (**e**), suggesting this incorrect detection was due to assembly error caused by segmental duplication formed by two homology genes, *GYPB* and *GYPA*.

**Fig. 6.**
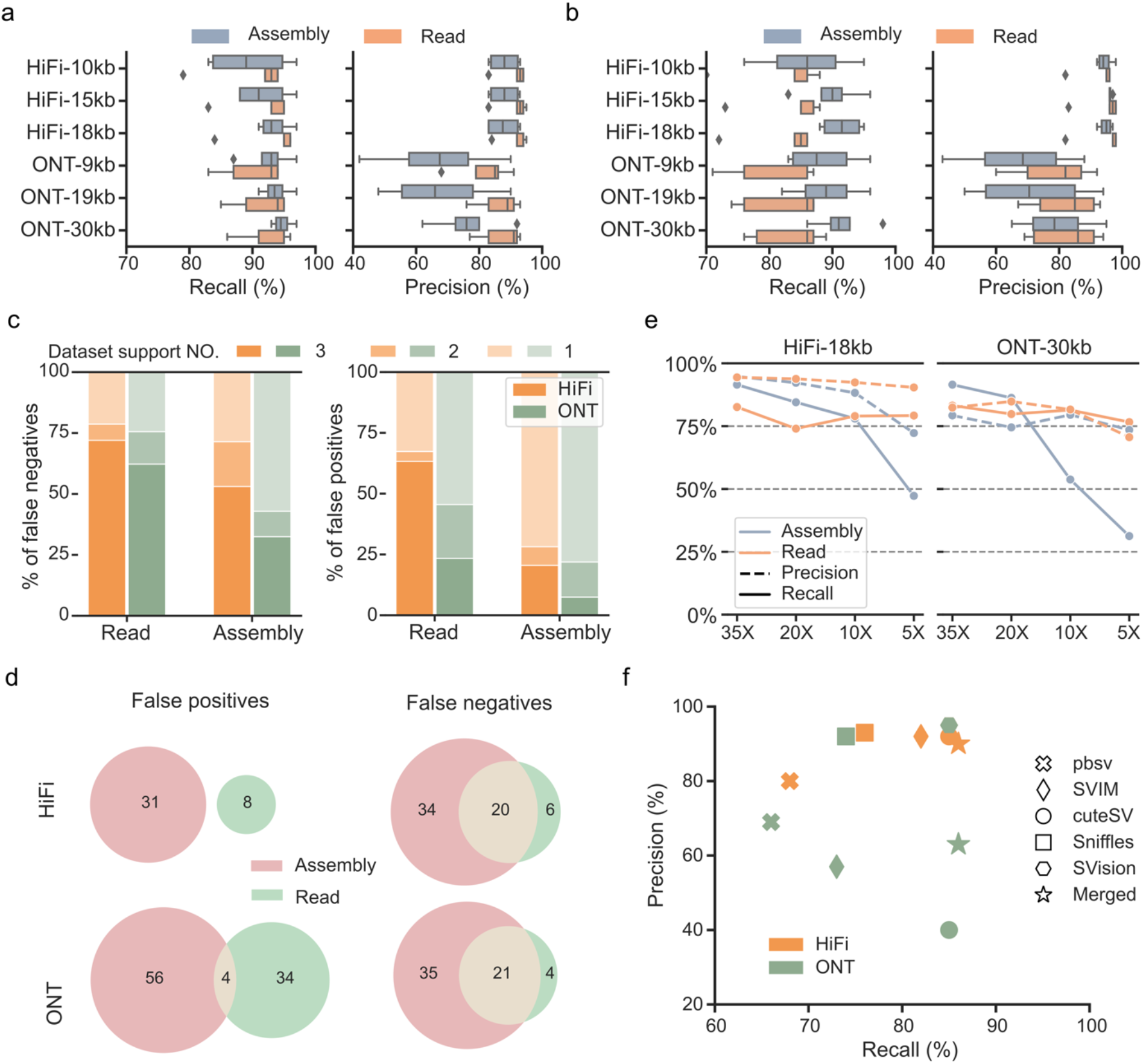
Summaries of benchmarking two strategies with well curated structural variants. **a**. The recall and precision of detecting structural variants (SVs) at true INS/DEL regions. **b**. The recall and precision of detecting SVs at challenging medically relevant autosomal genes (CMRGs). **c**. For SVs at CMRGs, percentage of false positive and false negative SVs among HiFi and ONT datasets, i.e., SVs in three, two and one dataset. **d**. The Venn-diagram of false positive and false negatives detected by both strategies on HiFi and ONT datasets. **e**. The impact of sequencing coverage on the recall and precision of detecting SVs at CMRGs. **f**. At 5X coverage, the recall and precision of each read-based callers as well as the merged callset.

### Impact of different settings on the reproducible SVs between strategies

The above analysis on each strategy suggested that read-based strategy was more versatile to different sequencing settings when reads mapped by the same aligner, while assembly-based strategy was less affected by assembler and its breakpoint was more accurate than read-based strategy on HiFi datasets. We then want to examine the impact of detection and sequencing settings on the reproducible SVs between strategies. In general, SVs were compared at whole genome scale and at 12,745 true insertions/deletions (INS/DEL) regions identified by GIAB [26]. Considering the number of used aligners, assemblers and callers, we obtained 128 merged sets of nonredundant SVs between strategies among six datasets. For the merged SV callsets, a median of 28,630 and 35,701 SVs at whole genome scale were identified, and a median of 14,141 and 15,840 SVs at true INS/DEL regions were identified from HiFi and ONT datasets, respectively (**Fig. 4a**). The unexpected large number of nonredundant SVs from ONT datasets were mainly contributed by merging PAV’s and SVision’s callsets (**Fig. 4b**).

Based on the nonredundant SV sets, we first assessed the impact of pairs of aligner and assembler on the number of concordant SVs between strategies, referring to as strategy concordant SVs. On average, 55% and 45% of the SVs at whole genome scale were strategy concordant SVs when detected with HiFi and ONT reads, respectively, and strategy concordant SVs took around 80% (HiFi datasets) and 70% (ONT datasets) of the SVs at true INS/DEL regions (**Fig. 4c, Supplementary Fig. 7a**). Remarkably, the highest concordant rate was 89% for SVs at true INS/DEL regions and 72% for SVs at whole genome scale, which was around 20% higher than SVs detected from ONT datasets (**Fig. 4c**). Moreover, using HiFi reads, we observed minor effect of assemblers on the average concordant SV rate but large variance caused by aligners. In particular, the concordant rate from highest to lowest was achieved by pairing with minimap2, winnowmap, lra and ngmlr without assembler bias (**Fig. 4c**), indicating the sequence alignment strategies of ngmlr and minimap2 were significantly different. Additionally, at whole genome scale, we observed a positive correlation between read length and strategy concordant SV rate on both HiFi and ONT datasets (**Fig. 4d**), and this correlation was expected because assemblers essentially created longer DNA sequences which equals to the usage of longer reads for SV detection. Afterwards, we examined the breakpoint consistency of strategy concordant insertions and deletions (INS/DEL), which dominated the discoveries of both strategies. On average, 77% and 74% of the concordant insertions and deletions were BSD-10 events when detected from HiFi and ONT dataset, respectively **(Fig. 4e**). However, we observed great platform bias for BSD-0 INS/DEL, where 38% of the insertions and 45% of the deletions were BSD-0 in HiFi callsets and it was around 20% higher than the percentage of BSD-0 INS/DEL detected with ONT reads (**Fig. 4e**). Furthermore, for breakpoint inaccurately reproduced SVs, 50% of the insertions and 83% of the deletions were found at simple repeat regions (**Supplementary Fig. 7b**).

To further understand the impact of assembler and aligner on the BSD-10 INS/DEL, we used BSD-10 INS/DEL detected from HiFi-18kb dataset because the highest concordant SV rate was observed on this dataset (**Fig. 4d**). Overall, detecting from minimap2 aligned reads, two strategies were able to detect the highest percentage of BSD-10 INS/DEL without assembler bias, and similar results were observed on winnowmap but significantly differed from ngmlr and lra (**Fig. 4f**). Especially for ngmlr, the highest percentage of BSD-10 INS/DEL was found between pbsv and any assembly-based callers without affecting by assembler (**Fig. 4f**). This was also consistent with our observation of BSD-10 INS/DEL among all datasets, where minimap2 and winnowmap performed similar but outliers were found among conordant SVs detected from ngmlr aligned reads (**Supplementary Fig. 7c**). Therefore, we reasoned that though 70% of the SVs were reproducible by both strategies and it was even 20% higher for SVs at true INS/DEL regions, further optimization of detecting SVs at complex genomic regions, especially tandem repeats, was required for future methods development.

### Examining SVs only detected by assembly-based strategy

Recently, several studies had claimed that assembly-based strategy is able to comprehensively detect SVs from an individual genome [3, 19]. Thus, we examined whether assembly only SVs (i.e., SVs only detected by assembly-based strategy but missed by all read-based callers) were also detectable by read-based strategy. Since the above analysis suggested that using longer reads mapped with minimap2 resulted in the fewest number of strategy unique SVs (**Fig. 4d, Supplementary Fig. 8a**), HiFi-18kb and ONT-30kb were used to assess the assembly only SVs (**Fig. 5a**). As a result, 4,265 assembly only SVs (1,630 and 2,635 SVs from HiFi and ONT datasets, respectively), consisting of 2,800 insertions and 1,465 deletions, were identified from HiFi-18kb and ONT-30kb datasets and most of them were heterozygous SVs (**Supplementary Fig. 8b**). Moreover, 77% of the assembly only SVs (74% on ONT and 81% on HiFi) overlapped with Simple Repeats, but around 25% of the SVs detected from ONT assemblies were found at Segment Dup regions (**Supplementary Fig. 8c**).

To examine whether 4,265 assembly only SVs were detectable from read alignments, we first noticed that most of these SVs were located at high mapping quality regions (average read mapping quality >= 20) (**Fig. 5b, Supplementary Fig. 8d**). Afterwards, we found that 64% (1,056 out of 1,630) and 51% (1,345 out of 2,635) of the assembly only SVs contain at least five HiFi and ONT SV signature reads identified from minimap2 alignments, respectively (**Fig. 5b**). These loci contain SV signature reads but missed by read-callers was mainly due to the large signature start position standard deviation, making them difficult to cluster for a valid call (**Fig. 5c**). Moreover, most of the average signature SV size ranged from 100bp to 1,000bp, which was not consistent with the size distribution of assembly only SVs at high mapping quality regions, especially for SVs smaller than 100bp (**Fig. 5c**). Therefore, even these assembly only SVs were reported by read-based callers, they were hard to match one event in assembly only SVs due to the breakpoint difference and size similarity. For those SV loci without enough SV signature reads, 65% (HiFi dataset) and 48% (ONT dataset) of the assembly unique calls overlapped with Simple Repeats (**Fig. 5d**). Additionally, on ONT dataset, 41% of the SVs without signature reads, consisting of 261 insertions and 182 deletions, overlapped with segmental duplications, which was six times than that on HiFi dataset (**Fig. 5d**). For example, an insertion of length 2,474bp (chr4:144,924,382-144,926,856) was detected from ONT assemblies at gene *GYPB* but no SV signatures found in HiFi read alignment and HiFi assembly alignment (**Fig. 5e**). Further investigation shows that gene *GYPB* had 97% sequence homology with *GYPA*, thereby leading to false discovery originated from assembly error (**Fig. 5f**). We also found an incorrect deletion of length 981bp at gene *SMPD4* without evident SV signature observed in HiFi reads and assemblies (**Supplementary Fig. 8e**). This gene was usually activated by DNA damage, cellular stress and tumor necrosis factor[27], and SVs associated with this gene had been identified in developmental disorder [28]. Therefore, we reasoned that read-based orthogonal validation is important and necessary to screen potential false discoveries from assembly-based calls, especially for clinical applications.

### Benchmarking strategies with SVs at complex genomic regions

The above analysis revealed that complex genomic regions, especially tandem repeat regions were hotspots for discordant SVs. To further assess SV detection performance, we used well curated HG002 SV at true INS/DEL regions and 203 SVs on CMRGs to evaluate two strategies, where SVs at true INS/DEL regions and CMRGs enabled the evaluation of SV detection at simple and complex genomic regions, respectively.

For the true INS/DEL regions, the highest recall was 97%, achieving by assembly-based strategy on both HiFi and ONT datasets, while the highest precision was achieved by read-based callers (**Fig. 6a**). Moreover, we noticed that the recall was positively correlated with read length for both strategies on HiFi and ONT datasets, but both strategies showed large precision variance on ONT datasets, especially for assembly-based strategy (**Fig. 6a**). As for SVs on CMRGs, assembly-based strategy outperformed the read-based strategy (**Fig. 6b**). Specifically, the highest recall of assembly-based strategy was 96%, and it was 7% higher than the highest one achieved by read-based strategy (**Fig. 6b**). Most importantly, we only noticed the positive correlation between recall and read length for assembly-based strategy without dataset preference (**Fig. 6b**). Furthermore, we investigated the false negative discoveries (i.e., missed benchmark SV) that affect recall of each strategy. As a result, 71% (54/76, HiFi) and 58% of (56/96, ONT) SVs detected by read-based strategy were false negative in three datasets, and these SVs were termed as datasets negatives, while the percentage of dataset negatives was 53% (26/49, HiFi) and 32% (25/77, ONT) for assembly-based strategy (**Fig. 6c, Supplementary Fig. 9a**). Similar to the above analysis, 63% of the false positive SVs (i.e., novel SVs detected by caller) detected by read-based strategy were concordant SVs among three HiFi datasets, i.e., referring to as datasets positives, which was 40% higher than assembly-based strategy on HiFi datasets (**Fig. 6c, Supplementary Fig. 9b**). The low of assembly-based strategy was due to the large number of false positive SVs detected from ONT-9kb dataset, i.e., 235 false positive SVs that were not found in dataset ONT-19kb and ONT-30kb (**Supplementary Fig. 9b**). We next compared the datasets negative and datasets positive SVs between two strategies, where two strategies tend to detected more concordant false negatives but false positives were often found to be strategy specifics (**Fig. 6d**).

Additionally, CMRGs are well documented across multiple diseases but often excluded from standard targeted or whole-genome sequencing analysis [26], enabling the evaluation for potential clinical application. The above analysis used the 35X coverage datasets, requiring around $7,000 and $3,000 for generating the HiFi and ONT reads, respectively, which was not applicable to clinical settings due to the high sequencing cost. Therefore, we subsampled the 35X coverage datasets to 5X, 10X and 20X coverage and examined the performance of each strategy. Overall, read-based strategy outperformed assembly-based strategy on both HiFi and ONT datasets when the coverage was below 20X (**Fig. 6e**). Remarkably, read-based strategy was sensitive when detected with 5X ultra-low coverage data, i.e., the average recall of read-based strategy was 78% for both datasets, and SVision and cuteSV achieved the highest recall and precision at such low coverage (**Supplementary Fig. 9c**). Moreover, the recall and precision of merged read-based callsets was slightly improved comparing to single caller while using 5X coverage data (**Fig. 6f**), which was consistent with other studies. At such low coverage, the average recall of assembly-based strategy was around 48% and 26% on HiFi-18kb and ONT-30kb dataset, respectively (**Fig. 6e**). Further investigation revealed that the low recall on ONT-30kb dataset was caused by assemblers, of which, the recall of calling SV from flye and shasta was 52% and 10%, respectively. However, such recall bias caused by assemblers on ONT dataset was not observed when detected from data of sufficient coverage, i.e., more than 20X (**Supplementary Fig. 9d**). The above results suggested that assembly-based strategy required at least 20X coverage data to achieve high recall and precision, but read-based strategy was able to achieve higher recall and precision with ultra-low coverage data, making it applicable to clinical screening.

## Discussion

Ongoing significant technology improvements have paved the way to apply long-read sequencing to population-scale sequencing projects and even for rapid genetic diagnoses, while the selection of proper SV detection strategy remains unclear. In this study, we compared and investigated the impact of factors that influenced the most widely-used read-based and assembly-based SV detection strategies. This is an important step towards the in-depth understanding of the usability and stability of each strategy in detecting SVs at genomic regions of different complexity as well as their potential application in clinical diagnosis.

For each strategy, we were able to identify the source of variability among different sequencing settings based on six long-read datasets. Our results showed that read-based strategy was versatile to different sequencing platforms once identical aligner was used, but applying assembly-based strategy on ONT datasets was greatly affected by assembler when compared to HiFi datasets. Notably, calling from HiFi assemblies, around 90% of the SVs could be reproduced among different datasets and it was slightly affected by assembler. Though flye was not comparable with hifiasm, it was flexible to both HiFi and ONT datasets and averagely 75% of the SVs were reproduced. Additionally, assembly-based strategy was able to identify more consistent breakpoint than read-based strategy for concordant SVs. We further investigated the impact of aligners and assemblers on each strategy. In terms of the number of reproducible SVs and their breakpoint consistency, SVs detected by assembly-based strategy were less affected by the usage of assemblers on HiFi datasets. On the contrary, concordant SV numbers, breakpoints and types of read-based callers were greatly affected by aligners, especially for ngmlr. Furthermore, we found that 70% of the whole genome scale SVs and 90% of the true INS/DEL region SVs were able to be detected by both strategies when proper assembler and aligner were paired. Most importantly, our results revealed a positive correlation between concordant SV rate and read length, incorporating with the recent achievements in generating reads of 4Mbp and longer [29], the percentage of reproducible is expected to be even higher. Furthermore, once considering assembly-based calls as a comprehensive callset, our analysis revealed that 66% and 52% of the assembly-based strategy uniquely detected SVs were detectable with read-based strategy on HiFi and ONT datasets, respectively, while they were missed because of the clustering issues caused by the signature ambiguity. This observation provided an important hint for future detection algorithm development.

The above comparison results provided supportive evidence of the strength and weakness of each strategy as well as the hotspots for discordant SVs. Accordingly, using well curated SVs at genomic regions of different complexity, we assessed the recall and precision of each strategy with different dataset settings. As a result, with sufficient sequencing coverage (at least 20X), assembly-based strategy outperformed read-based strategy for detecting SVs at true INS/DEL regions, especially for SVs at CMRGs. However, 20X coverage long-reads data is still not applicable to clinical applications due to the high sequencing cost. Further analysis with ultra-low coverage data (5X) revealed that read-based strategy is able to robustly detect SVs in challenging genes, where the sensitivity was even 30% higher than assembly-based strategy. Additionally, for low-coverage HiFi and ONT data, merging SVs from different callers slightly increased the sensitivity comparing to single callers, such as SVision and cuteSV, suggesting SV merge was no longer necessary for long-read based SV detection.

Moreover, our analysis showed that SVs at tandem repeat regions are the most challenging ones to detect consistently by two strategies, suggesting the demand of developing novel methods and data structures for resolving these SVs. These SVs are difficult to reproduce because calling from both read and assembly alignment can have systematic issues with misrepresented highly polymorphic loci in the linear reference genome, which only represent one allele and thus, do not incorporate repeat polymorphisms of a population [25]. To solve this issue, pan-genome reference, combing genomes from multiple individuals of a species, has been proposed improve SV detection at polymorphic regions as well as genotyping SVs using short-read data. Though graph methods offer great opportunity to solve bias for SV detection, these methods are still less straight-forward in practice then the use of linear reference genome. Moreover, it lacks evidence of how these graph-based methods generalize to clinical applications.

To the best of our knowledge, this was the first study of comparing the two representative long-read based SV detection strategies. Our analysis, from general-purpose detection to specific application, revealed the usability of each strategy, offering insights of selecting proper detection and sequencing settings for long-read projects. However, the evaluation is limited to diploid genomes and autosomal diseases, while the performance of two strategies on cancers, affecting by purity, heterogeneity and aneuploidy, requires further investigation.

## Conclusion

SV detection is an essential step for population genetics and clinical diagnosis. While a number of long-read based studies for both healthy and disease genomes had revealed the prominent performance of using read-based strategy and assembly-based strategy for SV detection, their strength and weakness toward different settings is yet to be assessed. In this study, systematic analysis of dataset concordant SV and strategy concordant SV revealed the impact of aligners, assemblers, read length and sequencing platforms on the usability and stability of two strategies, including breakpoint consistency and SV types. Afterwards, we have benchmarked each strategy on detecting SVs at genomic regions of different complexity, especially SVs at CMRGs. We expect this work will help users to select proper SV detection settings for different applications and foster future development of SV detection algorithms at complex genomic regions.

## Methods

### Read mapping and sequence assembly

The three HiFi datasets (i.e., HiFi-10kb, HiFi-15kb and HiFi-18kb) and the three ONT datasets (i.e., ONT-9kb, ONT-19kb, ONT-30kb) are all publicly available. Based on a recent review by Steve S. Ho et. al. [1], aligners containing minimap2, lra, winnowmap and ngmlr were included in our study, and assemblers including hifiasm, flye and shasta were used.

First of all, HiFi and ONT reads were mapped to human reference genome hg19 with minimap2 (v2.20), lra (v1.3.2), winnowmap (v2.03) and ngmlr (v0.2.7). Parameters used for each mapper were listed below:

- minimap2: parameters ‘*-a -H -k 19 -O 5,56 -E 4,1 -A 2 -B 5 -z 400,50 -r 2000 -g 5000*’ were applied to align HiFi reads, and ‘*-a -z 600,200 -x map-ont*’ were used for ONT reads.
- ngmlr: parameters ‘*-x pacbio*’ and ‘*-x ont*’ were used to align HiFi and ONT reads, respectively.
- winnowmap: parameters *‘-ax map-ont*’ and ‘*-ax map-pb*’ of winnowmap were used to map ONT and HiFi reads, respectively.
- lra: *‘-CCS*’ and ‘*-ONT*’ were set to map HiFi and ONT reads, respectively. We then applied each read-based caller with default parameters except the minimum number of SV supporting reads. Since the sequencing coverage was around 35X for all datasets, the minimum SV supporting read for each read-based caller was set to five for the detection of both homozygous and heterozygous SVs. For 5X coverage, the minimum SV supporting read for each read-based caller was set to one.

For sequence assembly, we use minimap2 aligned reads and phased SNPs released by GIAB to obtain phased reads via whatshap ‘haplotag’ option. Those unphased reads are randomly assigned as either haplotype 1 and haplotype 2, which are also used in further sequence assembly. Given the phased reads, we apply assemblers with default parameters to create the haplotype-aware assemblies.

### SV detection and post-processing

To detect SVs, methods were further excluded from the recent review [25] based on several criteria: (1) lack of detailed user manual; (2) no programming interface; (3) reported bias on aligners; (4) unresolved errors during wrapping. In the end, read-based callers including cuteSV (v1.0.10), pbsv (v2.2.2), SVIM (v1.4.0), Sniffles (v1.0.12) and SVision (v1.3.6) were selected and assembly-based callers including Phased Assembly Variant (PAV) and SVIM-asm were selected.

Read-based callers were directly applied to reads aligned by minimap2, ngmlr, lra and winnowmap with default parameters. Note that the minimum SV supporting read is set to five so that both homozygous and heterozygous germline SVs can be effectively detected from the 35X coverage datasets. For assembly-based strategy, the phased assemblies were directly used as input for PAV, and we run PAV with default parameters for SV detection. For SVIM-asm, assemblies were first mapped to reference hg19 with minimap2 parameters *‘-x asm20 -m 10000 -z 10000,50 -r 50000 --end-bonus=100 --secondary=no -O 5,56 -E 4,1 -B 5 -a*’, these parameters were used in minimap2 embedded in PAV. Then, we run SVIM-asm with parameters ‘*svim-asm diploid -- tandem_duplications_as_insertions --interspersed_duplications_as_insertions*’ for SV detection.

For each callsets, a BED file obtained from a publication [30] was used to exclude SVs located at centromere and other low mapping quality regions. SVs overlapped with regions in the BED file were ignored in the downstream analysis. For the rest of the autosome SVs, we then annotated their associated repetitive elements using Tandem Repeat Finder, RepeatMasker and Segmental Duplication results provided by UCSC Genome Browser. The original files downloaded from the genome browser were first processed based on scripts introduced by CAMPHOR [31]. Repeat element associated with each SV is assigned based on a recent publication [32]. In particular, Variable Number Tandem Repeat (VNTR) was assigned if the length of repeat unit longer than 7bp, otherwise, we considered it as Short Tandem Repeat (STR). It should be noted that simple repeat annotated by RepeatMasker was also classified into VNTR and STR. For SVs overlapping repetitive element, we require at least 50% of the entire SV length to be composed of the specific repeat type, and we prioritized the highest percentage of overlaps on the entire length of SV when multiple repeat types are annotated. For example, if 70% of an SV was composed of STR and 50% of the SV overlapped by ALU, then STR was assigned correspondingly. Moreover, according to the repetitive elements, we divided the genome into four different regions, i.e., Simple Repeat, Repeat Masked, Segment Dup and Unique. Simple Repeat represented regions of either VNTR or STR. Repeat Masked were those annotated as SINE, LINE, etc, by RepeatMasker. Segment Dup represented regions overlapping with segmental duplications. The rest of the genomic regions outside of Simple Repeat, Repeat Masked and Segment Dup were considered as Unique regions.

### Identification of concordant and unique SVs

According to different comparison purpose, we first obtained the nonredundant SVs of several callsets by running command ‘*Jasmine file_list=vcf_list*.*txt out_file=nonredundant_SVs*.*vcf max_dist=1000 spec_len=50 spec_reads=1’*. Then, using VCF file generated by Jasmine, we were able to identify concordant and unique calls as well as the breakpoint standard deviation of concordant calls. The breakpoint standard deviation was indicated in ‘STARTVARIANCE’ and ‘ENDVARIANCE’ in the VCF file. The major steps for analyzing SV reproducibility among datasets and strategies were listed as below:

- Dataset concordant/unique: Each caller was applied to six datasets for SV detection, and a nonredundant SV set was generated via Jasmine accordingly. SVs reproduced in six datasets were indicated by ‘SUPP=6’, while dataset unique calls were indicated by ‘SUPP=1’. Moreover, SVs reproduced by at least two datasets were indicated by ‘SUPP=2’, ‘SUPP=3’, ‘SUPP=4’, ‘SUPP=5’ and ‘SUPP=6’.
- Aligner concordant/unique: On each dataset, the reads were aligned with four aligners and SVs were detected subsequently with each caller. For a caller, we merged its four callsets originated from four aligners, from which, aligner concordant SVs were obtained with ‘SUPP=4’ and aligner unique SVs were labeled by ‘SUPP=1’.
- Assembler concordant/unique: On HiFi dataset, the reads were assembled by two assemblers (i.e., hifiasm, flye) and the assemblies were mapped with minimap2. For a caller, we merged its two callsets originated from two assemblers, from which, assembler concordant SVs were obtained with ‘SUPP=2’ and assembler unique SVs were labeled by ‘SUPP=1’. Similar process was applied to ONT dataset, but the assemblies were created by flye and shasta.
- Strategy concordant/unique: On each dataset, we obtained a nonredundant SV set between a read-based caller and an assembly-based caller via Jasmine. Strategy concordant and strategy unique calls were indicated by ‘SUPP=2’ and ‘SUPP=1’, respectively.

The breakpoint standard deviation of each SV in the merged set was kept in the ‘STARTVARIANCE’ column, and the values were directly used to analyze the breakpoint consistency of concordant SVs.

### Read alignment analysis for strategy unique calls

We applied the following steps to examine whether SVs uniquely detected by assembly-based strategy contain aberrant read alignment, i.e., the abnormal inter-read and intra-read alignments used to detect SVs by read-based callers.

- Step1. The assembly-based strategy uniquely detected SVs were classified to three types of regions according to the average read mapping quality (avg_mapq) obtained from minimap2 aligned reads:
  1. No read mapping region (No_reads)
  2. Low mapping quality regions (Low_mapq, avg_mapq < 20)
  3. high confident mapping regions (High_mapq, avg_mapq ≥ 20).
  The average mapping quality threshold 20 was set according to the default minimum read quality used for SV detection.
- Step2. The potential SV signature reads of those assembly unique SVs at high confident mapping quality regions were identfied. In general, the ‘I’ and ‘D’ tags in the CIGAR string, and the primary reads and their supplementary were collected and used to identify deletion (DEL), insertion (INS), inversion (INV) and duplication (DUP) signatures. The total number of reads containing SV signature was referred to signature count. Moreover, we calculated the start position standard deviation and size standard deviation of all signature reads.

### Evaluating each strategy with well curated SVs

For 35X coverage datasets HiFi-18kb and ONT-30kb, we down-sample them to 5X, 10X and 20X with SAMtools. Afterwards, each caller is applied to the 5X, 10X and 20X datasets with default parameters except for the number of minimum SV supporting reads, which is set to 1, 2 and 5 for 5X, 10X and 20X datasets, respectively. These values are set to enable effective detection of both homozygous and heterozygous germline SVs. The final VCF files are sorted, compressed and indexed for further evaluation. Furthermore, two benchmarks released by GIAB were used to assess both strategies of detecting SVs at true INS/DEL regions and CMRGs. The recall and precision were measured by Truvari with parameters ‘*-p 0*.*00 -r 1000 --passonly --giabreport*’, but the genotype accuracy was not considered in our evaluation.

## Availability of code and data

All related commands, analysis scripts and data download links are available at https://github.com/jiadong324/CompareStra.

## Ethics approval and consent to participate

Not applicable

## Author’s contributions

KY conceptualized and supervised the study. JL led the data analysis and conducted all performance comparison. SW contributed to structural variant analysis. PJ contributed to the sequence assembly. JL wrote the manuscript with input from all authors. All authors read and approved the final manuscript.

## Acknowledgements

The authors thank the comments from colleagues in the lab and those from HGSVC.

