## Supplementary Figures for "Comparison and benchmark of long-read based structural variant detection strategies"

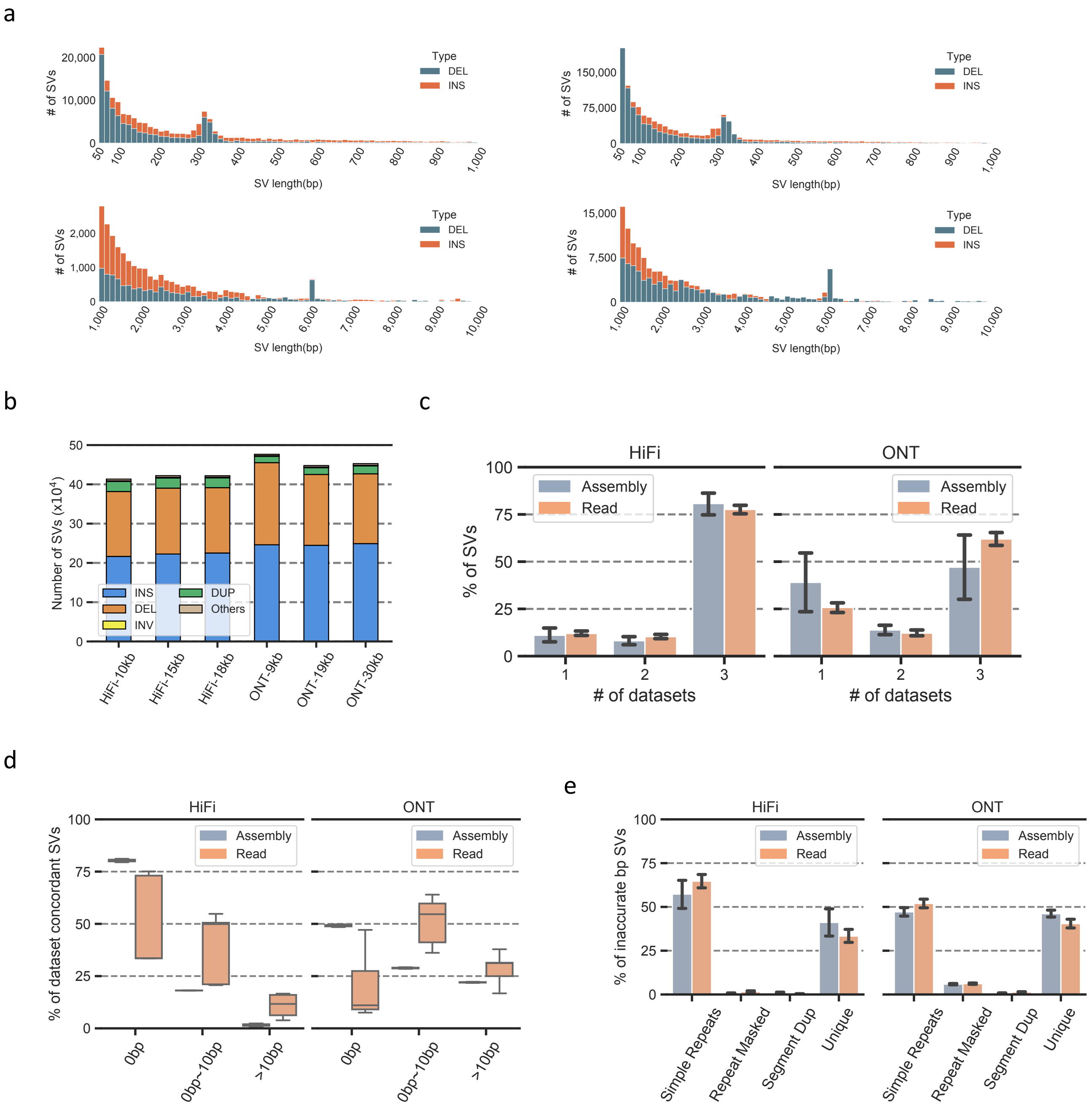


**Supplementary Fig. 1 a**. Size distribution of insertions and deletions detected by assembly-based (left) and read-based strategy (right). **b**. Number of different SV types detected by read-based strategy. **c**. Dataset concordant SVs of each strategy assessed on HiFi and ONT datasets separately. **d**. Breakpoint standard deviation of dataset concordant SVs detected by each strategy. **e**. Genomic regions of breakpoint inaccurately reproduced SVs, i.e., breakpoint standard deviation greater than 10bp.

a

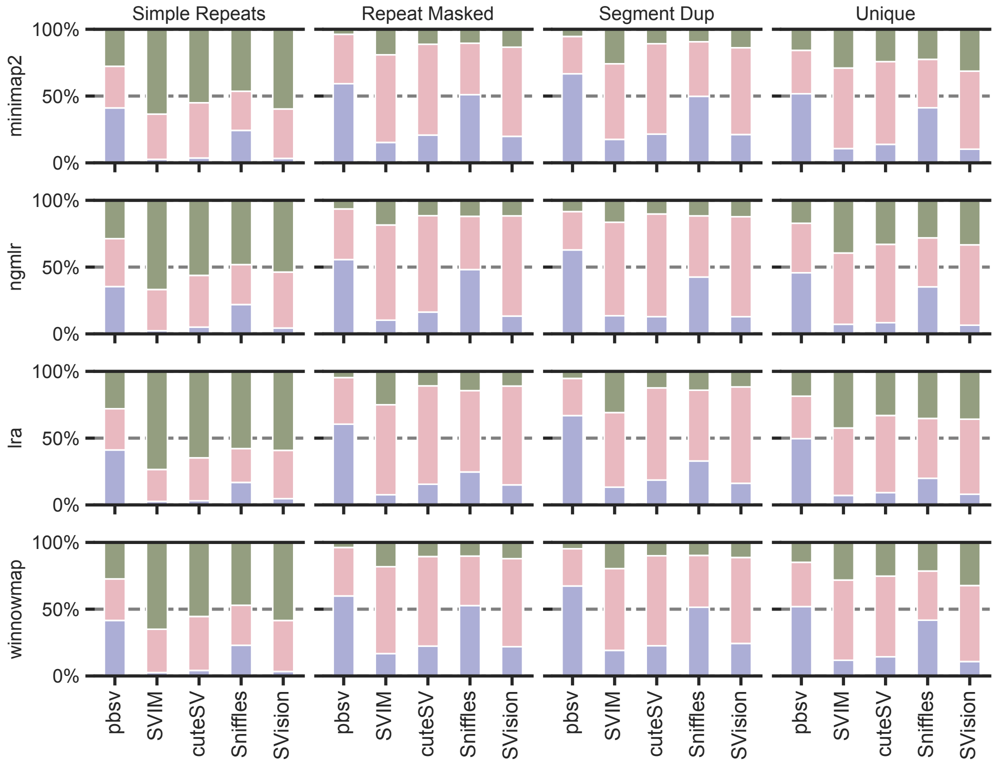


b

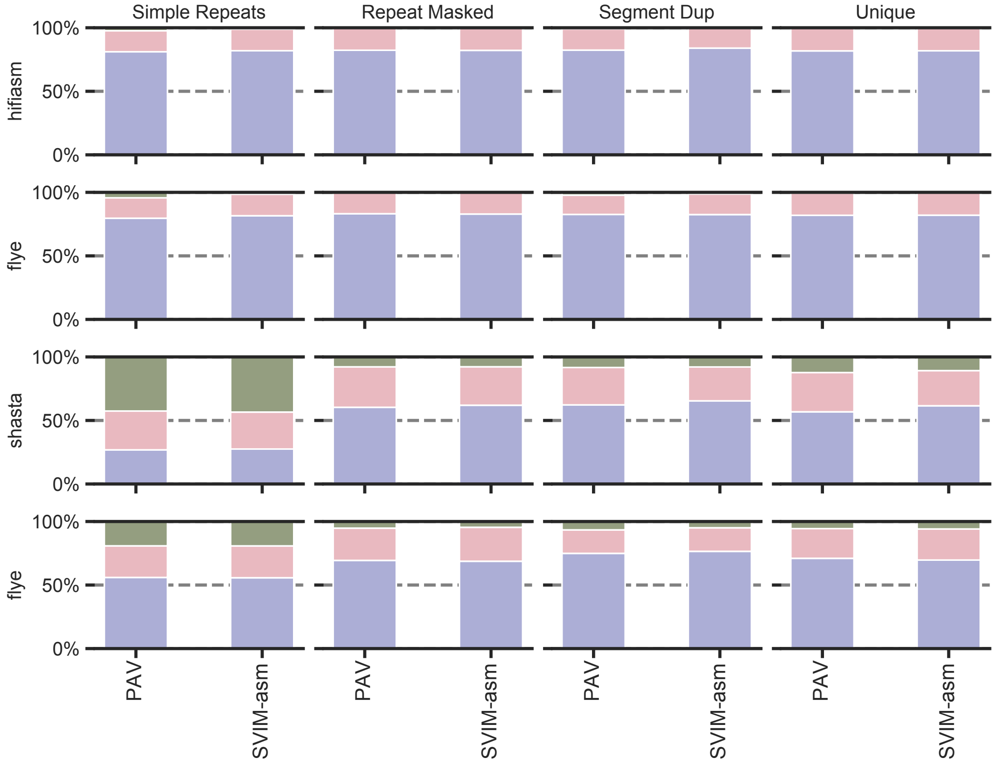


**Supplementary Fig. 2 a**. Breakpoint standard deviation of dataset concordant SVs detected by read-based callers on reads mapped with different aligners. **b**. Breakpoint standard deviation of dataset concordant SVs detected by assembly-based callers on assemblies created by different assemblers.

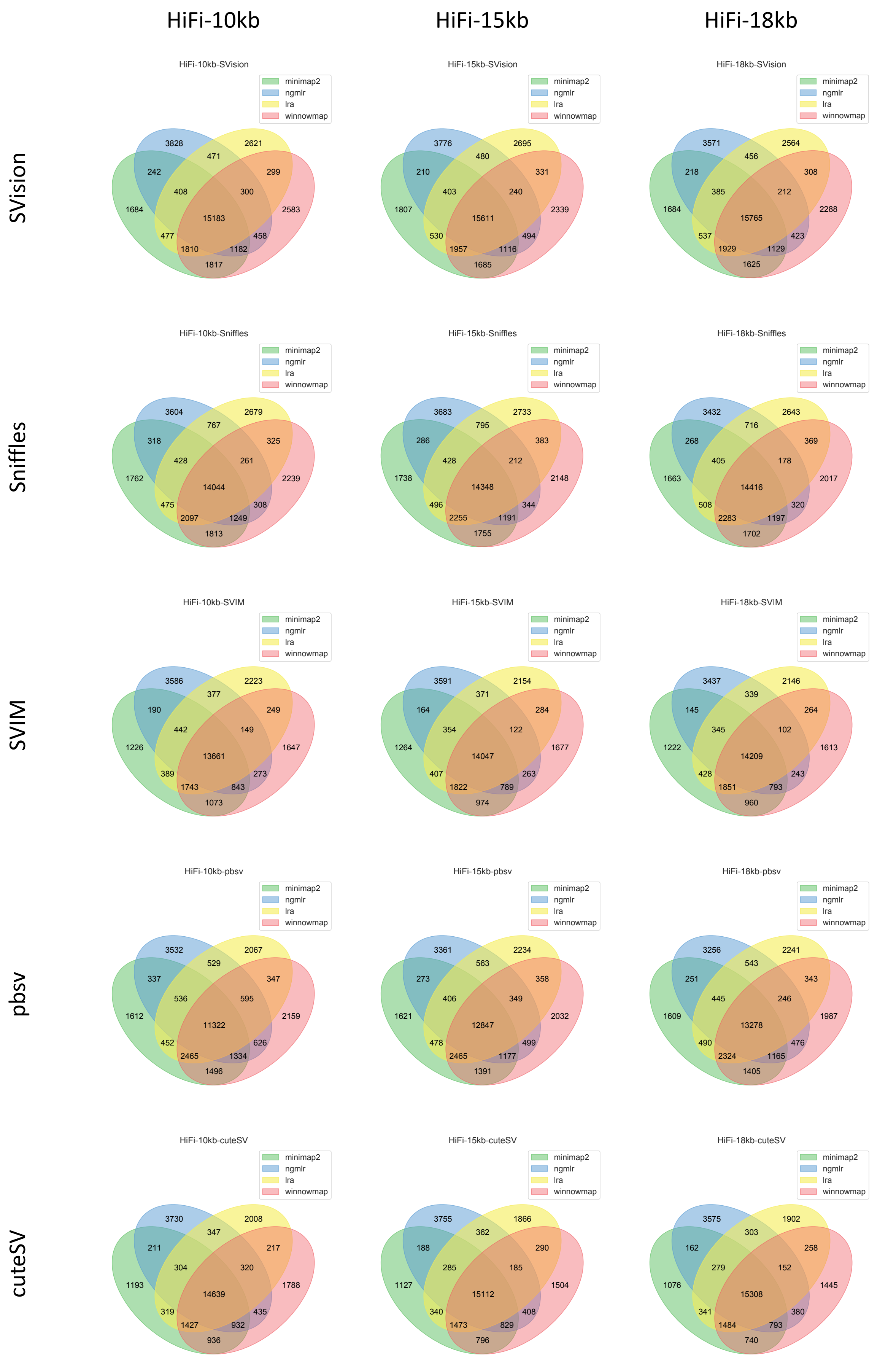


**Supplementary Fig. 3** Venn-diagram of aligner concordant SVs for each caller on HiFi datasets

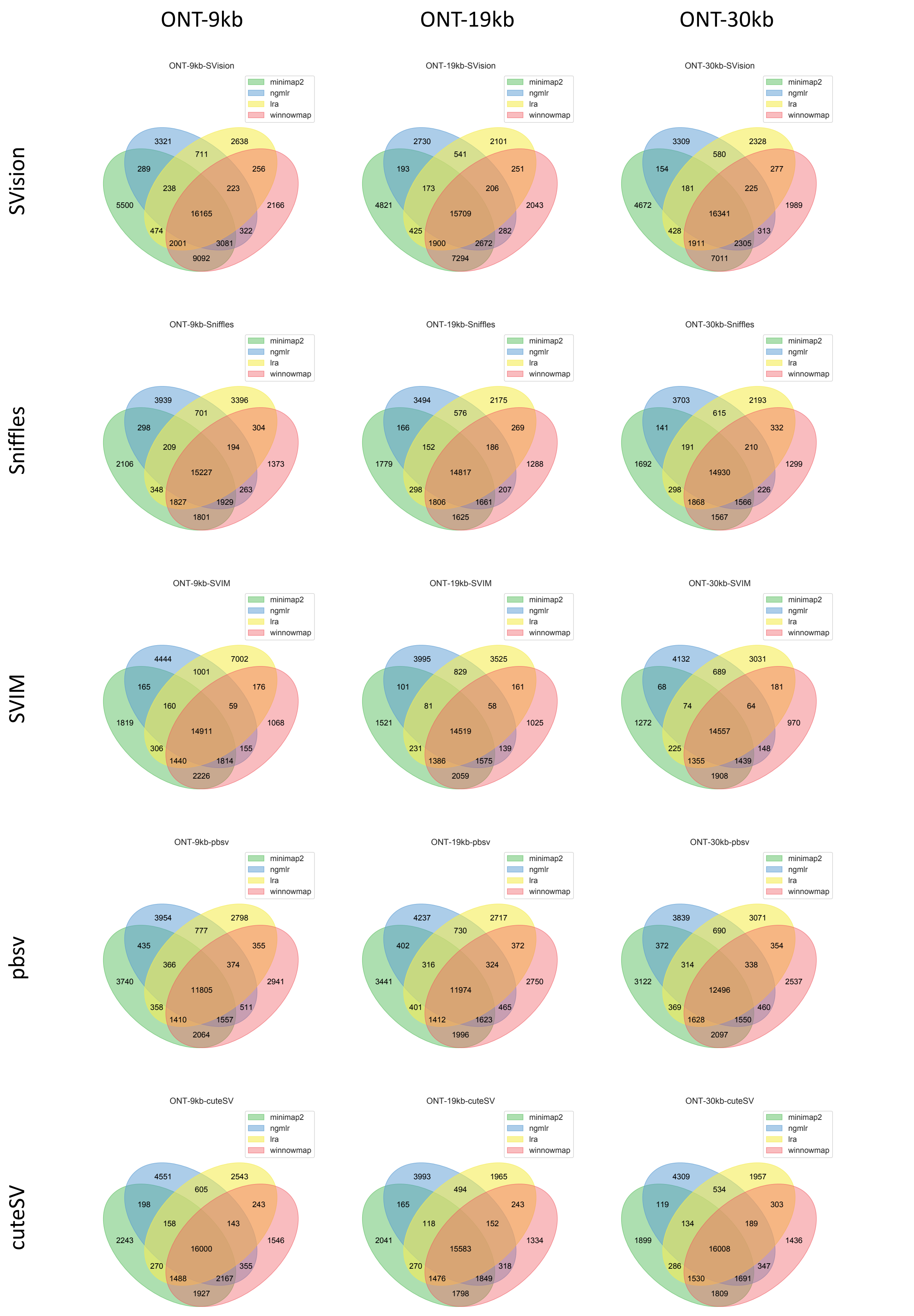


**Supplementary Fig. 4** Venn-diagram of aligner concordant SVs for each caller on ONT datasets

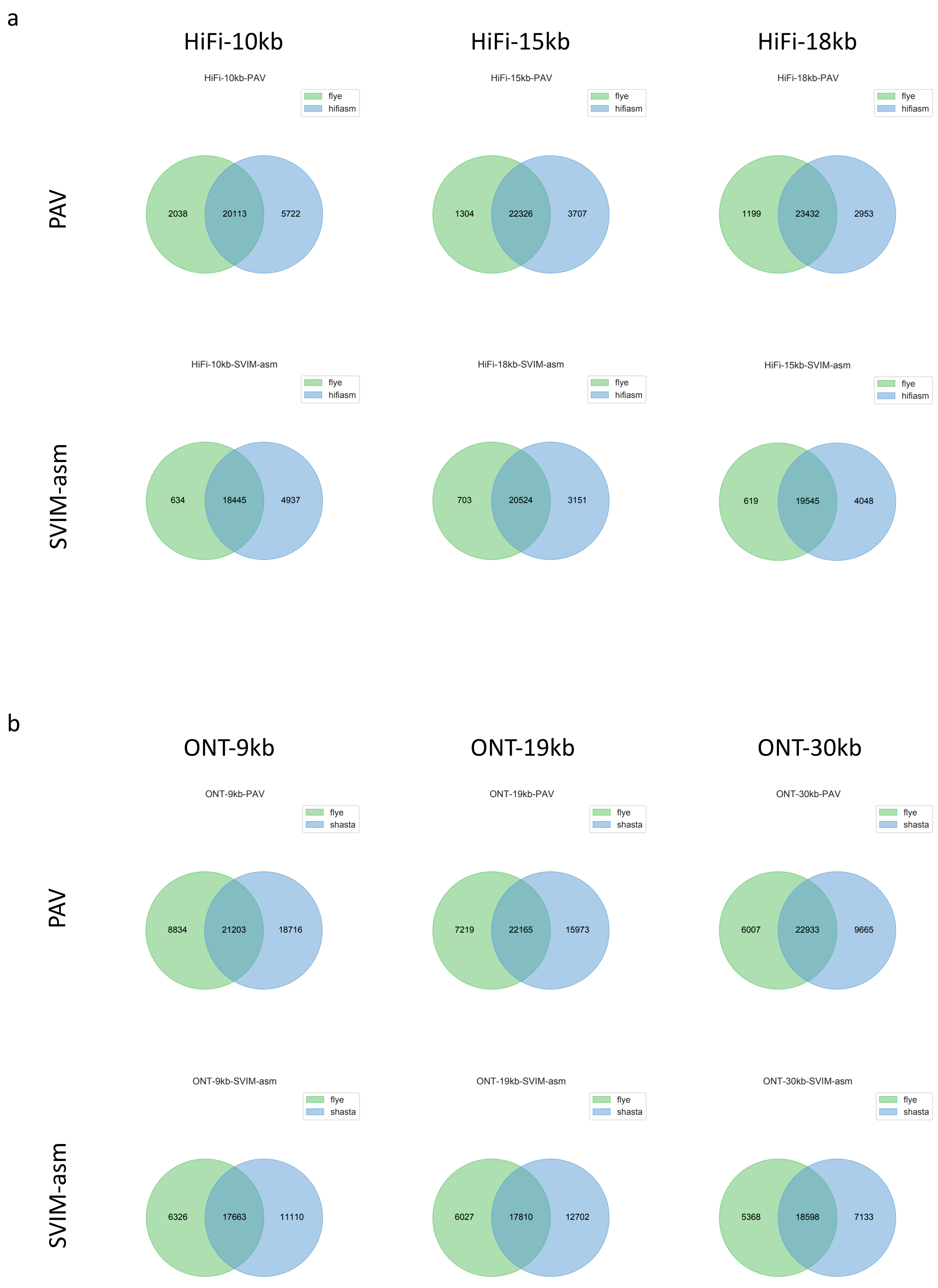


**Supplementary Fig. 5 a**. Venn-diagram of assembler concordant SVs for each caller on HiFi datasets. **b**. Venn-diagram of assembler concordant SVs for each caller on ONT datasets.

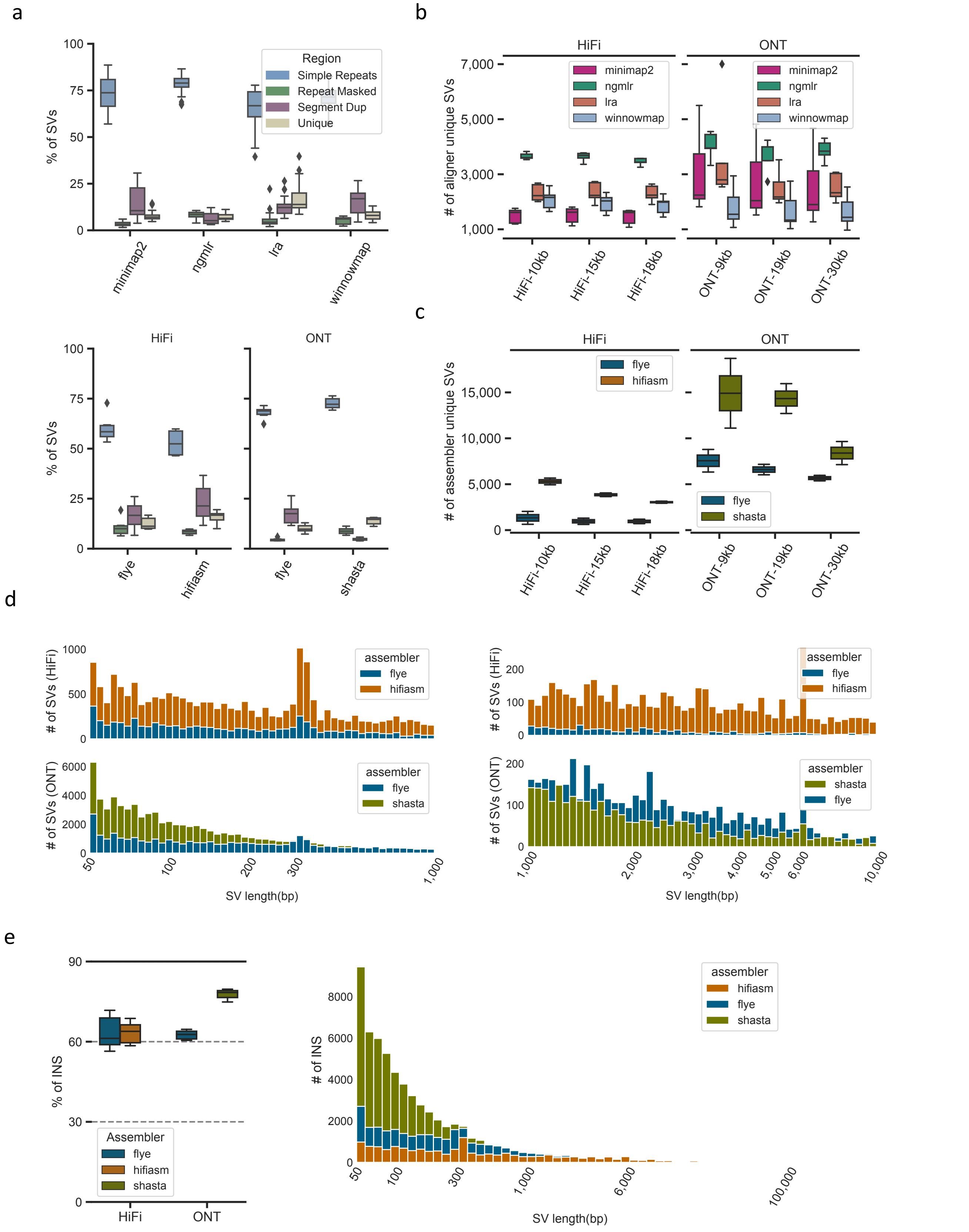


**Supplementary Fig. 6 a**. Genomic region distribution of aligner (top) and assembler (bottom) uniquely detected SVs. **b**. Distribution of the number of SVs uniquely detected from one of the aligners among different datasets. **c**. Distribution of the number of SVs uniquely detected from one of the assemblers among different datasets. **d**. Size distribution of SVs uniquely detected from one of the assemblers. **e**. Percentage insertions in SVs uniquely detected from one of the assemblers and their size distribution.

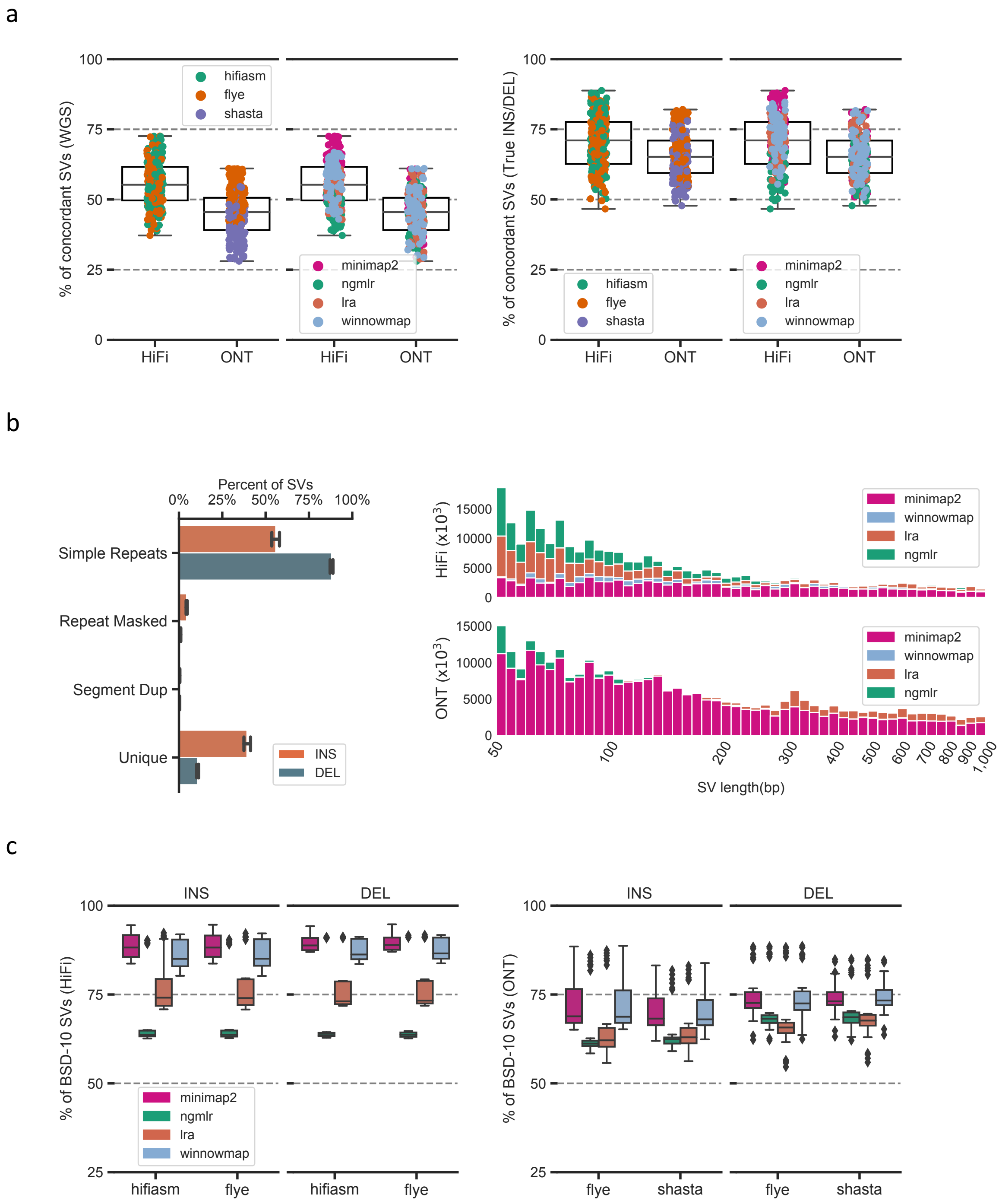


**Supplementary Fig. 7 a**. Percentage of SVs detected by both strategies at whole genome scale (WGS) and True INS/DEL regions. **b**. Genomic region distribution of breakpoint inaccurately reproduced strategy concordant SVs (left) and their size distribution (right). **c**. Percentage of breakpoint accurately reproduced strategy concordant SVs (i.e., BSD-10) affected by different aligner and assembler combinations.

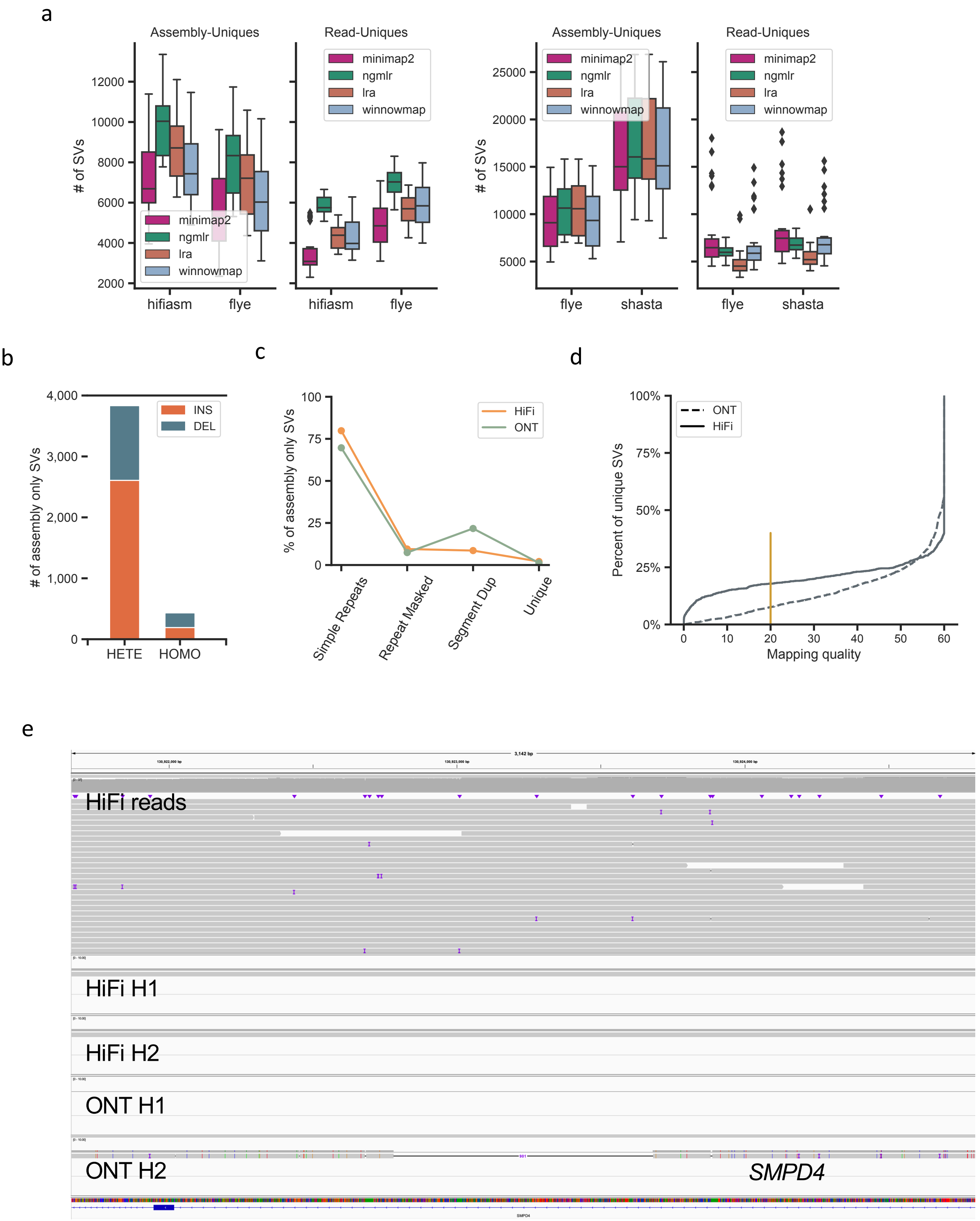


**Supplementary Fig. 8 a**. Number of SVs uniquely detected strategies affected by the aligner and assembler combinations. **b.** Number of heterozygous (HETE) and homozygous (HOMO) assembly only SVs. **c**. Distribution of assembly only SVs at genomic regions. **d**. Average mapping quality of assembly only SV loci. **e**. IGV view of an incorrect deletion at *SMPD4* detected by assembly-based strategy from ONT assemblies. From top to bottom tracks shows the alignments of HiFi read, HiFi haplotype-1 (H1), HiFi haplotype-2 (H2), ONT haplotype-1 (H1), and ONT haplotype-2 (H2).

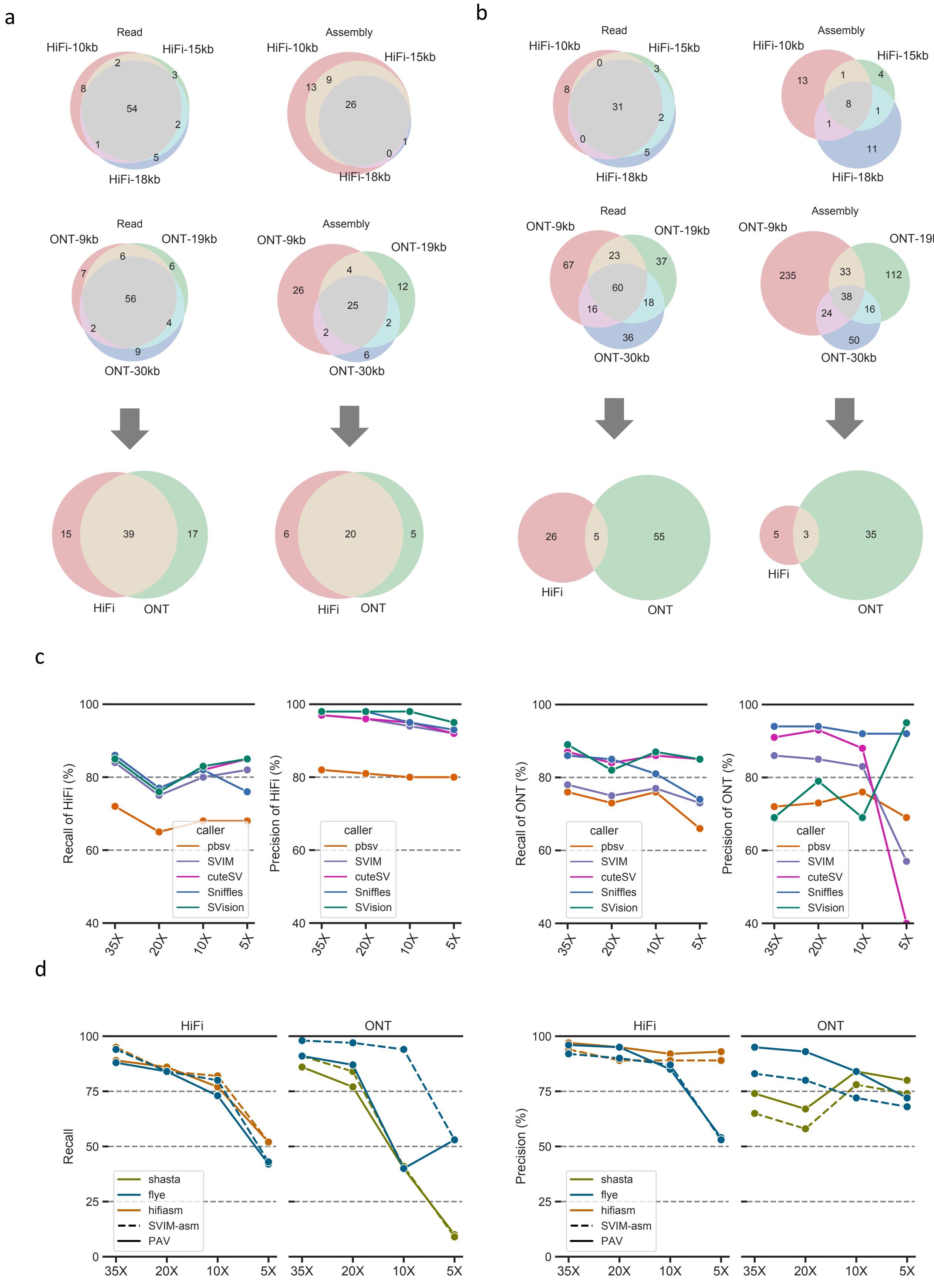


**Supplementary Fig. 9 a**. Venn-diagram of false negative discoveries of each strategy between datasets. **b**. Venn-diagram of false positive discoveries of each strategy between datasets. **c**. Recall and precision of detecting SVs on CMRGs with different coverage data. **d**. The impact of assemblers on the recall and precision of detecting SVs on CMRGs while using assembly-based callers at different coverages.

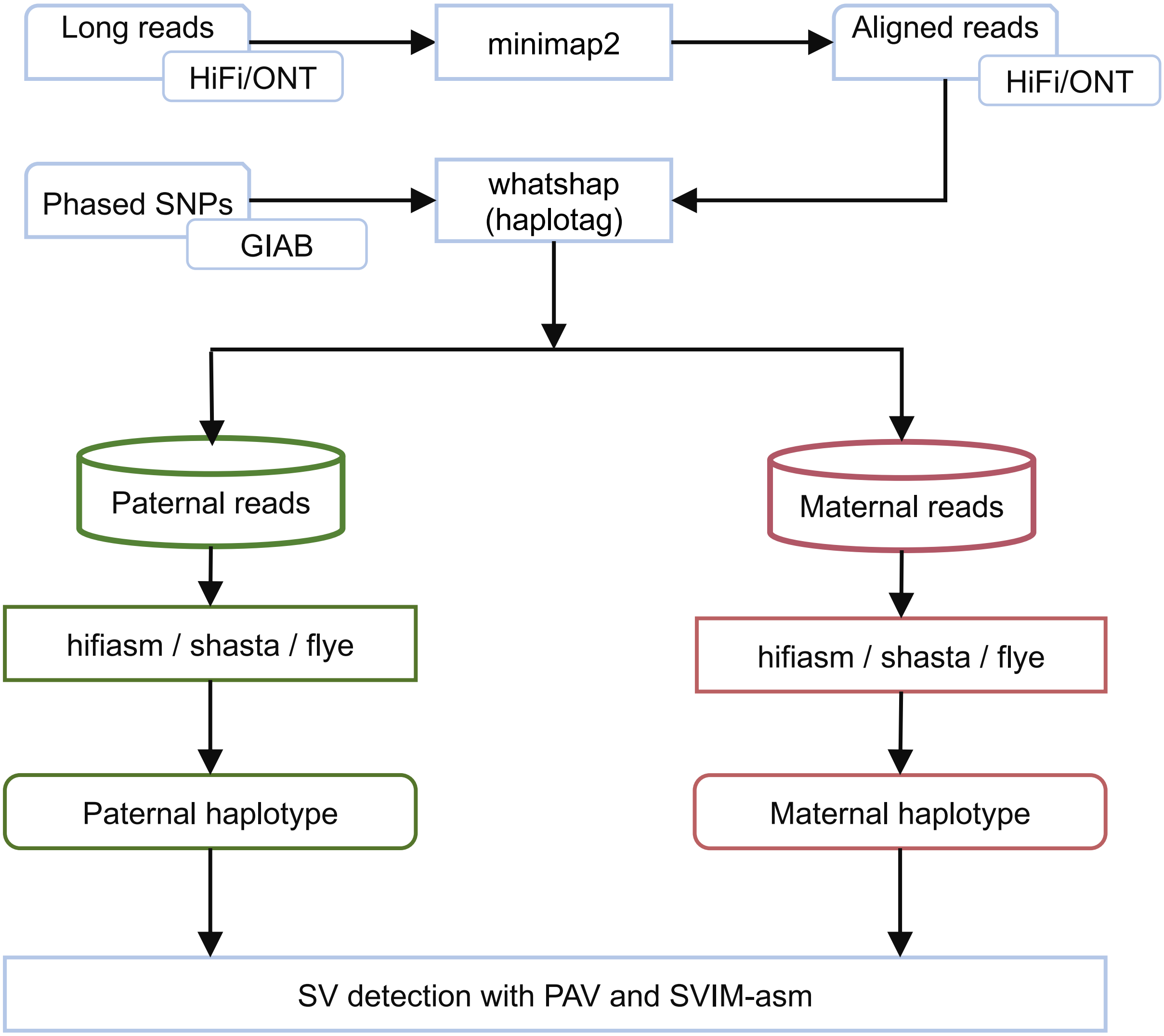


**Supplementary Fig. 10** Workflow of generating haplotype-aware assemblies
